# Cross-species transcriptomic analysis identifies mitochondrial dysregulation as a functional consequence of the schizophrenia-associated 3q29 deletion

**DOI:** 10.1101/2023.01.27.525748

**Authors:** Ryan H. Purcell, Esra Sefik, Erica Werner, Alexia T. King, Trenell J. Mosley, Megan E. Merritt-Garza, Pankaj Chopra, Zachary T. McEachin, Sridhar Karne, Nisha Raj, Brandon J. Vaglio, Dylan Sullivan, Bonnie L. Firestein, Kedamawit Tilahun, Maxine I. Robinette, Stephen T. Warren, Zhexing Wen, Victor Faundez, Steven A. Sloan, Gary J. Bassell, Jennifer G. Mulle

## Abstract

Recent advances in the genetics of schizophrenia (SCZ) have identified rare variants that confer high disease risk, including a 1.6 Mb deletion at chromosome 3q29 with a staggeringly large effect size (O.R. > 40). Understanding the impact of the 3q29 deletion (3q29Del) on the developing CNS may therefore lead to insights about the pathobiology of schizophrenia. To gain clues about the molecular and cellular perturbations caused by the 3q29 deletion, we interrogated transcriptomic effects in two experimental model systems with complementary advantages: isogenic human forebrain cortical organoids and isocortex from the 3q29Del mouse model. We first created isogenic lines by engineering the full 3q29Del into an induced pluripotent stem cell line from a neurotypical individual. We profiled transcriptomes from isogenic cortical organoids that were aged for 2 months and 12 months, as well as day p7 perinatal mouse isocortex, all at single cell resolution. Differential expression analysis by genotype in each cell-type cluster revealed that more than half of the differentially expressed genes identified in mouse cortex were also differentially expressed in human cortical organoids, and strong correlations were observed in mouse-human differential gene expression across most major cell-types. We systematically filtered differentially expressed genes to identify changes occurring in both model systems. Pathway analysis on this filtered gene set implicated dysregulation of mitochondrial function and energy metabolism, although the direction of the effect was dependent on developmental timepoint. Transcriptomic changes were validated at the protein level by analysis of oxidative phosphorylation protein complexes in mouse brain tissue. Assays of mitochondrial function in human heterologous cells further confirmed robust mitochondrial dysregulation in 3q29Del cells, and these effects are partially recapitulated by ablation of the 3q29Del gene *PAK2*. Taken together these data indicate that metabolic disruption is associated with 3q29Del and is conserved across species. These results converge with data from other rare SCZ-associated variants as well as idiopathic schizophrenia, suggesting that mitochondrial dysfunction may be a significant but overlooked contributing factor to the development of psychotic disorders. This cross-species scRNA-seq analysis of the SCZ-associated 3q29 deletion reveals that this copy number variant may produce early and persistent changes in cellular metabolism that are relevant to human neurodevelopment.

## Introduction

Rare variants have now been identified that confer extraordinarily high risk for schizophrenia (SCZ). Functional study of these variants may yield insights into the molecular and cellular impairments that ultimately give rise to psychosis. By restricting investigation to a single variant, etiologic heterogeneity is vastly reduced, which may lead to better discrimination of causal mechanisms. To date, the strongest identified single genetic risk factor for SCZ is the 3q29 deletion (3q29Del), a copy number variant (CNV) that encompasses 22 protein-coding genes and is located near the telomeric end of human chromosome 3 (Mulle et al., 2010; Mulle et al., 2016). Hemizygous loss of this set of genes is associated with at least a 40-fold increase in risk for SCZ (Marshall et al., 2017; Mulle, 2015); this deletion also increases risk for additional neurodevelopmental and psychiatric conditions, including intellectual disability, ASD, and ADHD (OMIM #609425) (Sanchez Russo et al., 2021).

Exciting developments in molecular neuroscience have led to extraordinary new tools for the investigation of neurobiology of mental health disorders. In this study, we leverage two state-of-the-art experimental model systems, which together amplify the rigor of our approach. The starting substrate for these experiments are two CRISPR-engineered experimental systems: newly-generated isogenic human induced pluripotent stem (iPS) cells, where we have precisely introduced the 3q29 deletion using CRIPSR/Cas9, and the 3q29 mouse model (B6.Del16^+/*Bdh1-Tfrc*^), which bears complete synteny to the human 3q29 interval and displays neurodevelopmental and somatic correlates of human syndromic phenotypes (Baba et al., 2019; Pollak et al., 2022; Rutkowski et al., 2019). These experimental systems offer complementary advantages; cortical organoids are the current gold standard model of human cortical development *in vitro,* whereas the syntenic 3q29Del mice provide a source of brain tissue from a physiological context. We hypothesized that a transcriptomic analysis of differentially expressed genes in developing cortical tissue would provide relatively unbiased insight into underlying mechanisms of cellular dysfunction. We reasoned that transcriptomic changes that are observed in *both* model systems are likely attributable to the 3q29Del and may underlie core phenotypes.

To investigate the biological effects of the 3q29Del, we performed single-cell mRNA sequencing (scRNA-seq) in isogenic human cortical organoids at both early (2-month) and late (12-month) developmental time points and perinatal (p7) mouse isocortex. We devised a strategy to systematically identify the most salient transcriptomic effects, both globally and in specific neural cell-types, to identify likely phenotypes for functional analysis. This strategy led us to a dysregulated transcriptome linked to mitochondria, which displayed both early and prolonged changes that were subsequently supported by orthogonal analyses of protein expression and functional assays. These results point to mitochondria as a possible site of convergent biology downstream of discrete neurodevelopmental variants.

## Materials and Methods

### Cell Culture and Genome Engineering

Whole blood samples of 5-10 mL were collected in EDTA Vacutainer test tubes and processed for the isolation of erythroid progenitor cells (EPCs) using the Erythroid Progenitor Kit (StemCell Technologies). EPCs were reprogrammed using Sendai particles (CytoTune-iPS 2.0 Reprogramming kit, Invitrogen) and plated onto Matrigel coated six-well plates (Corning). Cultures were transitioned from erythroid expansion media to ReproTesR (StemCell Technologies) and then fed daily with ReproTesR until clones were isolated. iPSCs were maintained on Matrigel coated tissue culture plates with mTeSR Plus (StemCell Technologies).

Cell lines were characterized for stem cell markers by RT-PCR and immunocytochemistry after at least 10 passages in culture. Total RNA was isolated from each cell line with the RNeasy Plus Kit (Qiagen) according to manufacturer’s protocol. mRNA was reverse transcribed into cDNA using the High Capacity cDNA Synthesis Kit (Applied Biosystems). Expression of pluripotency genes *OCT4*, *SOX2*, *REX1* and *NANOG* was determined by RT-PCR. Sendai virus inactivity was confirmed using Sendai genome specific primers.

Isogenic 3q29Del iPSC and HEK cell lines were generated using the SCORE method (Tai et al., 2016). To identify low-copy repeat (LCR) target sequences, the reference sequence (hg38) between *TNK2 – TFRC* (centromeric) and *BDH1 – RUBCN* (telomeric) was downloaded and aligned in NCBI BLAST. A ∼20 Kb segment was found to be 97% identical and was searched for gRNA sequences using CHOPCHOP (https://chopchop.cbu.uib.no) (Labun et al., 2019). Three single gRNA sequences (IDT) that were predicted to each cut at a single site in both LCRs were identified and cloned into pSpCas9(BB)-2A-Puro (PX459) V2.0, which was a gift from Feng Zhang (Addgene plasmid #62988; http://n2t.net/addgene:62988; RRID: Addgene_62988) (Ran et al., 2013).

Single gRNA plasmids were transfected into a neurotypical control iPSC line (maintained in mTeSR or mTeSR+ (STEMCELL, Vancouver) on Matrigel (Corning)-coated plates using a reverse transfection method and Mirus TransIT-LT1 reagent (Mirus Bio, Madison, WI) and transfected cells were transiently selected for puromycin resistance. Genome cleavage efficiency for each gRNA was calculated using the GeneArt Genomic Cleavage Detection Kit (Thermo) and gRNA_2 (5’-CAGTCTTGGCTACATGACAA-3’, directed to -strand, hg38 chr3:195,996,820 - chr3:197,634,397) was found to be the most efficient with cleaved bands at the predicted sizes. Cells transfected with gRNA_2 were dissociated and cloned out by limiting dilution in mTeSR supplemented with 10% CloneR (STEMCELL). Putative clonal colonies were manually transferred to Matrigel-coated 24-well plates for expansion and screened for change in copy number of the 3q29Del locus gene *PAK2* (Hs03456434_cn) using TaqMan Copy Number Assays (Thermo). Three (of 100) clones showed an apparent loss of one copy of *PAK2* and were subsequently screened for loss of the 3q29 genes *TFRC* (Hs03499383_cn), *DLG1* (Hs04250494_cn), and *BDH1* (Hs03458594_cn) and for no change in copy number to external (non-deleted) 3q29 genes *TNK2* (Hs03499383_cn) and *RUBCN* (Hs03499806_cn) all referenced to *RNASEP* (Thermo #4401631). All cell lines retained normal karyotypes (WiCell, Madison, WI) and were free of mycoplasma contamination (LookOut, Sigma).

To generate 3q29Del HEK-293T cell lines, HEK cells were transfected with either empty px459 or px459+gRNA_1 (5’-TTAGATGTATGCCCCAGACG-3’, directed to the +strand) and screened and verified with TaqMan copy number assays as described above. *PAK2* was deleted from a control HEK-293T line as detailed above (*PAK2* gRNA 5’-TTTCGTATGATCCGGTCGCG-3’, directed to -strand). Clones were screened by Western blot (Anti-PAK2 Abcam ab76293 1:5000, RRID: AB_1524149) and confirmed by Sanger sequencing PCR-amplified gDNA. HEK cell lines were also negative for mycoplasma contamination.

### Genome-wide Optical Mapping

1.5E6 iPSCs were pelleted, washed with DPBS, and frozen at −80°C following aspiration of all visible supernatant. 750ng of DNA was labeled, stained, and homogenized using the DNA Labeling Kit-DLS (Bionano; 80005). Stained DNA was loaded onto the Saphyr chip G1.2 and the chip was scanned in order to image the labeled DNA using the Saphyr System. Structural variants were called relative to the reference genome (hg38) using Bionano Solve. Structural variants were compared to the parent (unedited) cell line using the Bionano Solve Variant Annotation Pipeline.

### Cortical Organoid Differentiation

Engineered isogenic 3q29Del iPSC lines and the unedited parent line, along with two additional clonal lines from the same donor, were expanded in mTeSR or mTeSR+ on Matrigel-coated plates. On DIV 0, colonies were gently released from plates in 0.35mg/ml Dispase according to an established protocol (Sloan et al., 2018). Floating colonies were re-suspended in mTeSR supplemented with 10uM Y-27632 (Reprocell, Beltsville, MD) in ultra-low attachment 10cm dishes (Corning). After 48hr, spheroids were transitioned to Neural Induction Medium (20% Knockout Serum Replacement, 1% Non-essential amino acids, 100U/mL Pen/Strep, 0.5% GlutaMAX, 0.1mM 2-mercaptoethanol in DMEM/F12 w/ HEPES), supplemented with 5uM Dorsomorphin and 10uM SB-431542 (added fresh) with daily media changes through DIV 6. On DIV 7, Neural Induction Medium was replaced with Neural Medium (Neurobasal-A with 2% B-27 w/o vitamin A, 1% GlutaMAX, 100U/mL Pen/Strep) supplemented with fresh EGF (20ng/ml, R&D Systems) and FGF (20ng/ml, R&D Systems) for daily media changes through day 16. From day 17-25, organoids were fed Neural Medium with EGF and FGF every two days. From day 26-42, Neural Medium was supplemented with BDNF (20ng/ml, R&D Systems) and NT-3 (20ng/ml, R&D Systems) every two days. From day 43 onwards, organoids were fed Neural Medium without supplements twice weekly.

### Mouse Genotyping and Maintenance

All animal experiments were performed under guidelines approved by the Emory University Institutional Animal Care and Use Committee. Mice were genotyped as described previously (Rutkowski et al., 2019) and noted as either Control (wild-type, C57BL/6 N Charles River Laboratories) or 3q29Del (B6.Del16^+/*Bdh1-Tfrc*^, MGI:6241487). Male 3q29Del mice and Control littermates were included in the scRNA-seq study. Both male and female mice were included in mitochondrial fractionation experiments.

### Tissue Dissociation and Sorting

Single-cell suspensions from cortical organoids (DIV 50 = “2-month” N=2 Control, N=2 3q29Del, and DIV 360 = “12-month” N=2 Control, N=2 3q29Del) and postnatal day 7 (P7) mouse cortices (N=4 Control, N=4 3q29Del) were produced by a papain dissociation method based on a published protocol (Foo, 2013). Tissue was coarsely chopped with a sterile scalpel and digested for 1hr at 34°C in a pH-equilibrated papain solution (Worthington, Lakewood, NJ) with constant CO_2_ flow over the enzyme solution. Digested tissue was gently spun out of papain, through ovomucoid solutions, and sequentially triturated with P1000 and P200 pipet tips. Live cells were counted by manual and automated methods (Countess II, Thermo) and in organoid samples were isolated from cellular debris by fluorescence-activated cell sorting on a FACSAria-II instrument (calcein AM-high, Ethidium Homodimer-1 low).

### Single-cell Library Prep and RNA-Sequencing

Single-cell suspensions were loaded into the 10X Genomics Controller chip for the Chromium Next GEM Single Cell 3’ kit workflow as instructed by the manufacturer with a goal capture of 10,000 cells per sample. The resulting 10X libraries were sequenced using Illumina chemistry. Mouse samples and libraries were prepared and sequenced at a separate time from human samples.

### scRNA-seq Data Processing and Analysis

To quantify gene expression at single-cell resolution, the standard Cell Ranger (10x Genomics) and Seurat (Satija et al., 2015) data processing pipelines were followed for demultiplexing base call files into FASTQ files, alignment of scRNA-seq reads to species-specific reference transcriptomes with STAR (mouse: mm10, human: GRCh38), cellular barcode and unique molecular identifier (UMI) counting, and gene- and cell-level quality control (QC). To filter out low-quality cells, empty droplets and multiplets, genes expressed in <10 cells, cells with >30% reads mapping to the mitochondrial genome, and cells with unique feature (gene) counts >7,000 were removed based on manual inspection of the distributions of each QC metric individually and jointly. Outlier cells with low unique feature counts were further removed via sample-specific thresholding of corresponding distributions (<250 for mice; <700 for organoids). Thresholds were set as permissive as possible to avoid filtering out viable cell populations, consistent with current best-practice recommendations (Luecken and Theis, 2019).

The *sctransform* function in Seurat was used for normalization and variance stabilization of raw UMI counts based on regularized negative binomial regression models of the count by cellular sequencing depth relationship for each gene, while controlling for mitochondrial mapping percentage as a confounding source of variation (Hafemeister and Satija, 2019). Resulting Pearson’s residuals were used to identify the most variable features in each dataset (n=3,000 by default), followed by dimensionality reduction by PCA and UMAP, shared nearest neighbor (SNN) graph construction on the basis of the Euclidean distance between cells in principal component space, and unbiased clustering of cells by Louvain modularity optimization. Optimal clustering solutions for each dataset was determined by building cluster trees and evaluating the SC3 stability index for every cluster iteratively at ten different clustering resolutions with the *clustree* function in R (Zappia and Oshlack, 2018). The effect of cell-cycle variation on clustering was examined by calculating and regressing out cell-cycle phase scores in a second iteration of *sctransform*, based on the expression of canonical G2/M and S phase markers (Tirosh et al., 2016). Consistent with the developmental context of the interrogated datasets, cell-cycle differences were found to covary with cell-type and retained in final analyses as biologically relevant sources of heterogeneity. Cluster compositions were checked to confirm comparable distributions of experimental batch, replicate ID, and genotype metadata. Cluster annotations for cell-type were determined based on the expression of known cell-type and cortical layer markers curated from the literature (Loo et al., 2019; Rosenberg et al., 2018; Ximerakis et al., 2019; Zhang et al., 2014; Zhang et al., 2016). Clusters exhibiting cell-type ambiguity were further sub-clustered to refine annotations or dropped from downstream analysis in case of inconclusive results (human cl.7 and cl.16; mouse cl. 25 and cl. 27).

Differential gene expression testing for genotype was performed on log normalized expression values (scale.factor=10,000) of each cluster separately with a two-part generalized linear model that parameterizes the bimodal expression distribution and stochastic dropout characteristic of scRNA-seq data, using the MAST algorithm, while controlling for cellular detection rate (Finak et al., 2015). A threshold of 0.1 was implemented as the minimum cut off for average log-fold change (logfc.threshold) and detection rates (min.pct) of each gene in either genotype to increase the stringency of differential expression analysis. Multiple hypothesis testing correction was applied conservatively using the Bonferroni method to reduce the likelihood of type 1 errors, based on the total number of genes in the dataset. To facilitate comparative transcriptomics, human homologs (including multiple paralogs) were identified for all differentially-expressed genes (DEGs) in the mouse dataset via the NCBI’s HomoloGene database (ncbi.nlm.nih.gov/homologene/). Data processing and analysis pipelines were harmonized across the mouse and organoid datasets, yielding parallel computational approaches for cross-species comparison of differential expression signals. The BrainSpan Developmental Transcriptome dataset used for developmental stage estimations was obtained by bulk RNA-Sequencing of postmortem human brain specimens collected from donors with no known history of neurological or psychiatric disorders, as described previously (BrainSpan, 2013; Kang et al., 2011). This large-scale resource is accessible via the Allen Brain Atlas data portal (https://www.brainspan.org/static/download/, file name: “RNA-Seq Gencode v10 summarized to genes”); dbGaP accession number: phs000755.v2.p1. All statistical analyses of scRNA-seq data were performed in R (v.4.0.3).

### scRNA-seq Pathway Analysis

To interpret differential gene expression results, pathways likely impacted by the 3q29Del were determined based on statistically over-represented gene-sets with known functions using g:Profiler (Raudvere et al., 2019). DEGs (Bonferroni adj. p<0.05) for each cluster were identified as described above and input with an experiment-specific background gene set (genes with min.pct > 0.1 in any cluster). GO:Biological Process (GO:BP) and Reactome (REAC) databases were searched with 10 < term size < 2000. Significantly enriched pathways below a threshold of g:SCS < 0.05 (Reimand et al., 2007) were compiled and filtered in Revigo (Supek et al., 2011) to reduce redundancy and determine umbrella terms.

### Seahorse Mitochondrial Stress Assay

HEK-293T cells that had been engineered to carry the 3q29Del (*3q29Del*), PAK2 knockout (*PAK2*), and mock-edited control cells (*CTRL*) were plated on poly-D-lysine coated 96-well Seahorse assay plates (XF96, Agilent) in DMEM (Gibco A144300) supplemented with 10% FBS, 2mM L-glutamine, 1mM sodium pyruvate, and either 10mM D-(+)-glucose (“Glu”, 7.5E3 cells/well) or 10mM galactose (“Gal”,15E3 cells/well). After 48hr, cells were washed twice in XF DMEM Assay Medium (Agilent) with either glucose or galactose (10mM) supplemented with 1mM pyruvate, 2mM glutamine.

Mitochondrial stress test compounds were loaded into injection ports as indicated by the manufacturer to achieve the following final concentrations: 1uM oligomycin, 0.25uM FCCP, 0.5uM rotenone, 0.5uM antimycin A (all sourced from Sigma). Cells equilibrated at 37°C with ambient CO_2_ for approximately 1hr prior to assay initiation. At the end of the experiment, cells were washed twice in PBS+Ca^2+^+Mg^2+^ and lysed at 4C for 30 min in 0.5% Triton X-100 protein buffer (150mM NaCl, 10mM HEPES, 0.1mM MgCl_2_, 1mM EGTA, 1x HALT Protease+Phosphatase inhibitor). Protein concentrations in each well were determined by BCA (Pierce) to normalize oxygen consumption rate data. Data were analyzed in Wave (Agilent). Four independent experiments were performed.

### Mouse Brain Mitochondrial Isolation

A protocol for mitochondrial isolation was adapted from prior work (Nagy and Delgado-Escueta, 1984). Two whole brains per genotype were dissected from adult mice (2-6mos.) and pooled in 2.5mL of ice cold Medium I (0.32M sucrose, 5mM HEPES pH 7.5, 0.1mM EDTA, Complete protease inhibitor) and homogenized with 16 strokes at approx. 800rpm in a Teflon glass homogenizer (0.125mm clearance) with a rest on ice mid-way through. Crude homogenate was cleared by centrifugation at 1000 x *g* for 10min and the supernatant was further centrifuged at 12,000 x *g* for 20min. All centrifugations were carried out at 4°C.

Isoosmotic Percoll (9 parts Percoll to 1 part 2.5M sucrose vol/vol) gradients were prepared in Medium II (0.25M sucrose, 5mM HEPES pH 7.2, 0.1mM EDTA). The second pellet was carefully re-suspended in an appropriate volume of 8.5% Percoll to produce a 7.5% Percoll solution and then was gently homogenized by twisting the Teflon pestle through the solution. The 7.5% Percoll solution containing the re-suspended tissue fraction was carefully layered on top of a gradient containing 16% and 10% Percoll. Gradients were centrifuged for 20min at 15,000 x *g* and mitochondrial fractions were extracted from the bottom of the tube and solubilized in 0.5% Triton X-100 protein buffer (150mM NaCl, 10mM HEPES, 0.1mM MgCl_2_, 1mM EGTA, 1x Complete). Protein concentrations were determined by BCA (Pierce) and normalized. 20ug of protein was loaded to each lane of Criterion gels for SDS-PAGE. Gels were transferred onto PVDF membranes by standard protocols and blocked in 5% milk. OXPHOS complex component proteins were probed for with an OXPHOS antibody cocktail (1:250, Abcam ab110412, RRID: AB_2847807). Protein levels were determined by band densitometry and quantified by normalizing to the most stable complex component (V).

## Results

### Generating isogenic 3q29Del iPSC lines

To isolate the effects of the 3q29Del from variable human genetic backgrounds, we introduced the full 1.6 Mb deletion into an iPSC line derived from a neurotypical individual by adapting a method previously used to generate isogenic iPSC lines carrying other neurodevelopmental CNVs (Tai et al., 2016). Like most recurrent CNVs, the 3q29Del is flanked by LCRs or segmental duplications (SDs), which are multi-kilobase stretches of highly homologous sequence (Bailey et al., 2002) that are likely involved in the formation of structural variants such as CNVs (Stankiewicz and Lupski, 2002). We targeted this homologous sequence with a single guide RNA that is predicted to cut at one site within each 3q29 SD (Labun et al., 2019) and isolated three clonal lines carrying the 3q29Del (Supp. Fig. 1). All clones retained normal iPSC morphology and karyotype, and genome-wide optical mapping analyses revealed no off-target structural variants.

### Single-cell transcriptomics in developing mouse and human cortical tissue

Three deletion clones and three clones of the parent cell line were differentiated to dorsal forebrain cortical organoids by established methods (Supp. Fig. 1f) (Sloan et al., 2018). Single-cell transcriptomes were produced from multiple organoids from two clonal lines per genotype at 2-months and 12-months into *in vitro* differentiation to capture a broad diversity of developing and mature cell-types. 54,255 cells were included in the human cortical organoid analysis (54% Control, Supp. Fig. 2). A mean of 2,805 genes were detected in each cell.

The 3q29Del mouse has been previously reported by two independent groups to express neuropsychiatric phenotypes including alterations in startle responses and social interactions (Baba et al., 2019; Rutkowski et al., 2019). Postnatal day 7 (P7) was chosen for tissue dissociation and single-cell sequencing to capture an array of mature and developing cell-types. 71,066 cells were included in the mouse scRNA-seq analysis (52.9% Control, Supp. Fig. 3). The mean number of genes detected in each cell was 2,920.

Fig. 1c and 1e show the major cell-types with distinct expression profiles that were isolated in each sequencing experiment. As expected, 2-month and 12-month human cortical organoids contained many of the cell-types that were also found in the perinatal mouse isocortex including excitatory neurons, astrocytes, immature neurons, radial glia/neural stem cells, neural progenitors, choroid plexus/ependymal cells, and oligodendrocyte progenitors (Supp. Fig. 4). In addition to these cells, we also found immune cells, inhibitory neurons, vascular cells, and endothelial cells in the mouse experiment (Supp. Fig. 5). Notably, in both experiments, we did not observe large-scale changes in cell clustering by genotype (Fig. 1d, 1f), but did observe a stark division in human cell clustering by time point (Fig. 1d). Indeed, most human clusters were comprised almost entirely of cells from a single time point, and only one cluster (cl. 2, annotated as migrating neuroblasts) was nearly evenly split by time point, consistent with developmentally regulated shifts in cell-type composition (Supp. Fig. 2). Predictably, astrocytes, oligodendrocyte progenitors, and upper layer excitatory neurons were not yet present in 2-month organoids (Sloan et al., 2018) but were indeed found in 12-month organoids.

**Fig 1.**
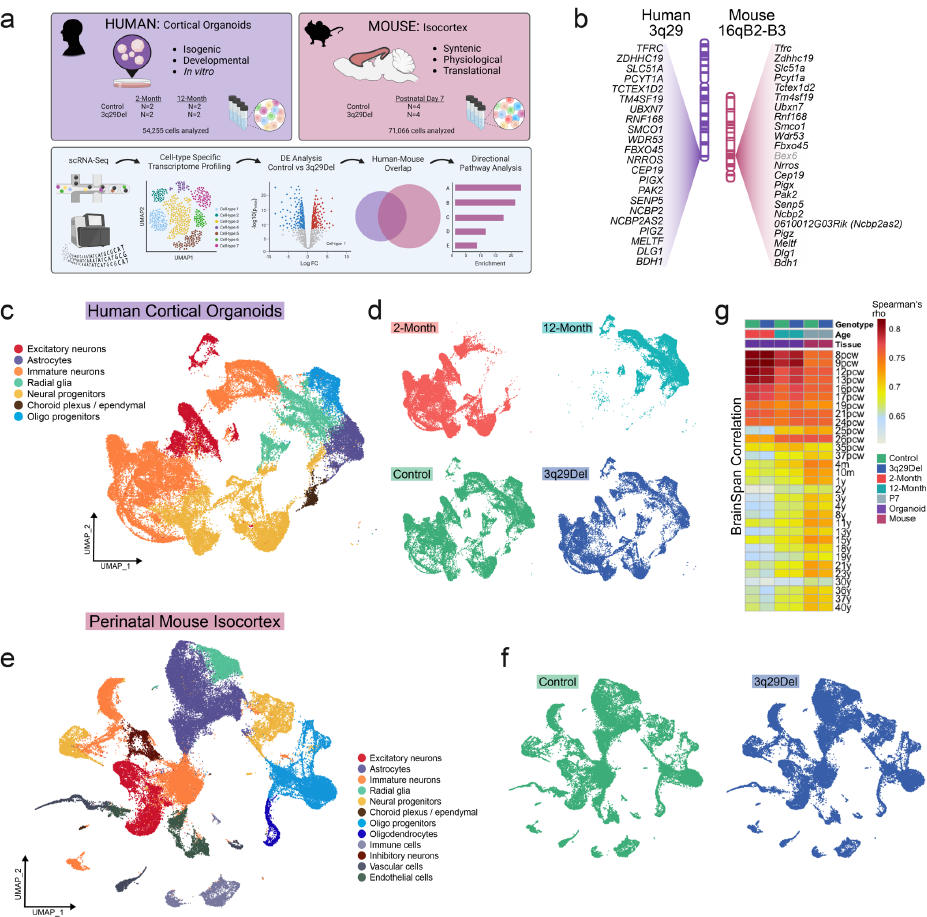
Cross-species single-cell sequencing. (**a**) A single-cell RNA-sequencing experiment was performed in isogenic human induced-pluripotent stem cell (iPSC)-derived cortical organoids at two time points and in postnatal day 7 mouse isocortex. An overview of the strategy to collect and filter differential gene expression data from both model systems is illustrated. (**b**) The human 3q29 deletion locus is nearly perfectly syntenic with a region of mouse chromosome 16, with the same gene order inverted. Corresponding loci are illustrated in the same orientation to facilitate clearer cross-species comparison. *Bex6* (in gray) is the only gene present in the mouse, not in the human locus. (**c** and **e**) UMAP dimensionality reduction plots colored by the main cell-types identified in human (**c**) and mouse (**e**) experiments. Human and mouse cells showed no obvious difference in gross distribution by genotype (**d**, **f**) but human cells were clearly divided in their transcriptomic clustering patterns by time point (**d**, top). The average expression profile of each sample was correlated (Spearman) to BrainSpan gene expression data profiling the human brain transcriptome in postmortem specimens across the lifespan (Kang et al., 2011) (**g**). Abbreviations: pcw, post-conception weeks (prenatal); m, months (postnatal); y, years (postnatal).

To better understand the window of cortical development that our experimental models best reproduce, we compared the average expression profile of each sample to postmortem human brain transcriptomes from the BrainSpan database (Kang et al., 2011). Spearman correlations revealed that 2-month organoids best matched very early phases of human brain development (8-9 post-conception weeks, Spearman π>0.80), whereas 12-month organoids and P7 mouse cortical cells maintained strong correlations through the second trimester of human gestation (π>0.76, Fig. 1g). As a control experiment, we compared the gene expression profiles of the human homologs of Control and 3q29Del mouse liver (Pollak et al., 2022) to the same BrainSpan data and, as expected, found that all correlations were markedly poorer than with any human or mouse cortical sample (π<0.56, Supp. Fig. 3). Together, these data suggest that the gene expression profiles of 2-12 month human cortical organoids and P7 mouse cortex best model the first two trimesters of human gestation.

We performed differential expression analysis in each cluster of both experiments by genotype. In both mouse and human cells, all 3q29Del transcripts were decreased to approximately match copy number in nearly every cell-type (Supp. Fig. 6-8). The only mouse-specific gene located in the syntenic locus (*Bex6*) was either not detected or not differentially expressed in any cluster. Across all clusters and two time points, there were 5,244 unique DEGs in human organoids, and 3,482 DEGs across all mouse clusters. To test for a similar global impact of the 3q29Del, all unique DEGs were compiled for human and mouse experiments. Strikingly, we found that more than half of the 3,253 strictly-matched human homologs of mouse DEGs were also found to be DEGs in the human dataset (Supp. Fig. 9a, fold enrichment = 1.53, hypergeometric p = 1.64E-162).

We explored whether the total number of DEGs in a given cluster was determined by cluster size (i.e. number of cells assigned to a cluster) or the number of 3q29 locus DEGs found in that cluster (Supp. Fig. 9b). In both mouse and human datasets, the number of 3q29 DEGs within the cluster (but not cluster size) was found to be a significant predictor of the total number of DEGs (negative binomial regression, p<0.0001), suggesting that haploinsufficiency of genes in the 3q29 locus is a significant driver of total differential gene expression.

To further test the degree of similarity in differential gene expression across mouse and human experiments, we plotted the average log fold change of DEGs in 10 comparable clusters and ran Pearson correlations (Supp. Fig. 9c-l). From proliferating neural progenitors to deep layer excitatory neurons, we found significant positive correlations (moderate to strong) in mouse and human 3q29Del gene expression changes in all comparisons except in astrocytes (Supp. Fig. 9h), which may reflect differences in maturation state between 12-month cortical organoid cells and postnatal mouse brain.

### Effects of 3q29Del on expression of mitochondrial and metabolic genes

We developed two systematic approaches to understand the most salient effects of the 3q29Del on the developing cortical transcriptome in mouse and human models. First, we sought to determine the pathways that were most frequently enriched across mouse and human clusters regardless of cell-type. To identify these frequently implicated pathways, DEGs from each cluster were split by direction of change (up-regulated vs down-regulated) and pathway analysis was performed as described in methods. All significantly enriched Gene Ontology: Biological Process (GO:BP) pathways were compiled and filtered by Revigo (Supek et al., 2011) to identify umbrella terms that were frequently dysregulated across multiple clusters.

We found that *Oxidative Phosphorylation* (OXPHOS) was both down-regulated and up-regulated across multiple clusters (Fig. 2b). A closer examination revealed that all down-regulated OXPHOS clusters were found in 12-month organoids and all up-regulated OXPHOS clusters were found in 2-month organoids. Moreover, the glycolysis-related *Pyruvate Metabolic Process* was found to be down-regulated in several of the 2-month clusters (cl.3, cl.11) that also up-regulated OXPHOS. This observation inspired the hypothesis that 3q29Del cells may exhibit altered metabolic maturation. A critical stage of neuronal differentiation is the switch from the heavily glycolytic progenitor state to mitochondrial aerobic respiration in mature neurons, which involves down-regulation of several key genes including *LDHA* (Zheng et al., 2016), which encodes the enzyme lactate dehydrogenase A. Cluster-level analysis revealed a striking decrease in the expression of *LDHA* in 3q29Del early-born deep layer excitatory neurons (cl. 3) and neural progenitors (cl. 6, cl. 11), which also showed increased mitochondrial gene expression (Fig. 2d). This result indicates a possible alteration in neuronal metabolic transition. Notably, all 12-month clusters displayed down-regulated OXPHOS (Fig. 2c, e), which may indicate a long-term effect of 3q29Del on cellular aerobic respiration across multiple neural cell-types.

**Fig 2.**
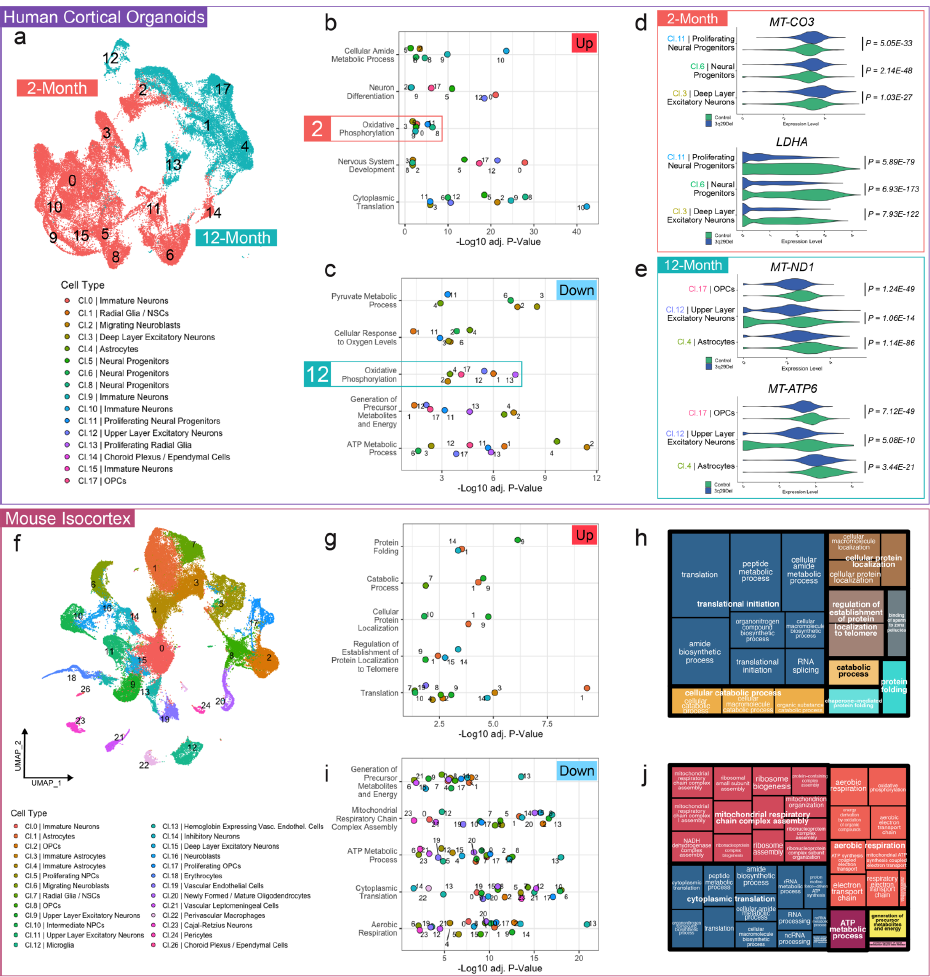
Transcriptomic evidence of metabolic changes in 3q29Del. The umbrella pathways most frequently found to be differentially-expressed based on up-(**b**) and down-(**c**) regulated genes in cortical organoids (**a**). Oxidative phosphorylation (OXPHOS) was enriched among both increased and decreased genes, but all clusters contributing to up-regulated OXPHOS were from 2-month organoids and all clusters contributing to down-regulated OXPHOS were from 12-month organoids. (**d**) Example violin plots visualizing log-normalized expression data of genes dysregulated in 2-month organoid clusters: *MT-CO3* (increased in 3q29Del) encodes the respiratory chain complex IV subunit COX3, *LDHA* (decreased in 3q29Del) is a key enzyme in glycolysis. (**e**) Example violin plots visualizing log-normalized expression data of genes dysregulated in 12-month organoid clusters: *MT-ND1* (decreased in 3q29Del) encodes a component of respiratory chain complex I and *MT-ATP6* (decreased in 3q29Del) encodes a component of the ATP Synthase complex. The most frequently up-(**g**) and down-(**i**) regulated umbrella pathways in mouse isocortex are shown. Treemaps derived by Revigo analysis (**h** and **j**) display the hierarchical organization of specific Gene Ontology Biological Processes (GO:BP) identified in pathway analysis. Similar colors denote semantic similarity. The size of each rectangle is proportional to the number of clusters exhibiting over-representation of a given GO:BP term. (All p-values are adjusted for multiple comparisons). Abbreviations: OPC, oligodendrocyte progenitor cells; NSC, neural stem cells; cl, cluster.

Differential pathway analysis of the mouse cortical data further supported the notion of a long-term decrease in mitochondria-related gene expression. Of the top 5 most frequently down-regulated umbrella terms, 4 were related to mitochondrial function and energy metabolism (Fig. 2i-j). Most mouse clusters were found to have down-regulated genes enriched for at least one of *Aerobic Respiration, ATP Metabolic Process, Mitochondrial Respiratory Chain Complex Assembly,* and *Generation of Precursor Metabolites and Energy*.

### Cross-species analysis in astrocytes and neurons

A second, parallel analysis strategy that we employed was to stringently filter exact-match DEGs in homologous human and mouse clusters by direction of change. First, we identified the human homologs of mouse DEGs in astrocytes (mm cl.1) and compared them to human DEGs in organoid cl. 4 (Fig. 3a). We found a 1.96-fold enrichment of commonly down-regulated genes (hypergeometric p = 5.51E-4) but no significant overlap among up-regulated DEGs (Fig. 3b). Pathway analysis revealed that the 29 commonly down-regulated genes were heavily enriched for terms related to the electron transport chain and OXPHOS (Fig. 3c).

**Fig 3.**
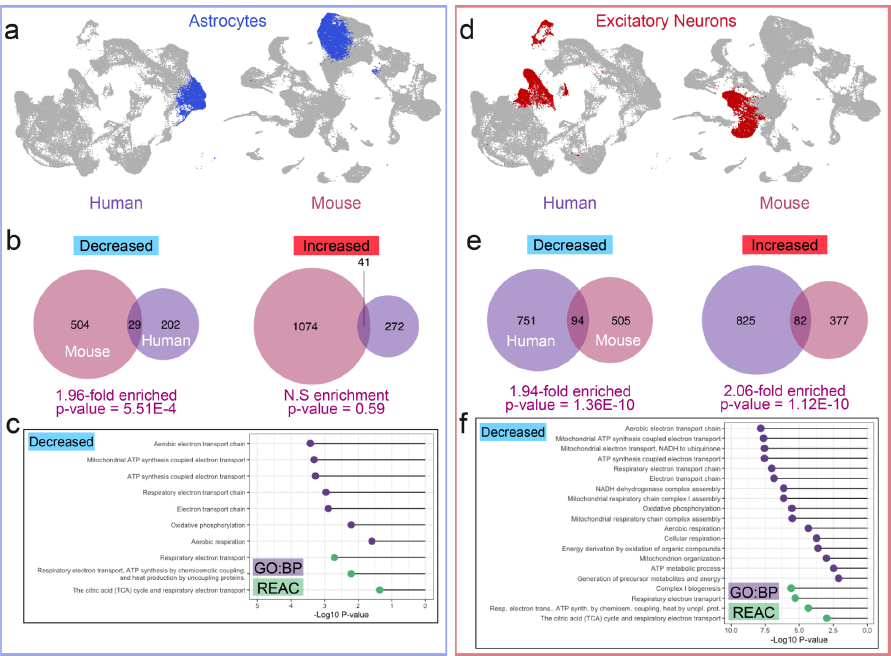
Common patterns of differential gene expression in two major mouse and human cell-types. Astrocytes were identified in human cortical organoids (12-month) and mouse isocortex. Corresponding clusters are color coded in blue in UMAP projections (**a**). The human homologs of mouse DEGs identified by MAST analysis were compared to organoid DEGs based on direction of change and a significant overlap was observed between the down-regulated DEGs of mouse and organoid astrocyte clusters (**b**). Pathway analysis of overlapping DEGs showed that all significantly enriched Gene Ontology: Biological Process (GO:BP) and Reactome (REAC) terms were related to mitochondrial function and metabolism (**c**). Upper and deep layer excitatory neuron DEGs were pooled and unique organoid DEGs were compared to the human homologs of mouse DEGs based on direction of change. Corresponding clusters are color coded in red in UMAP projections (**d**). There was a significant overlap between the DEGs of mouse and organoid excitatory neuron clusters for both up-regulated and down-regulated genes (**e**). Decreased genes were heavily enriched for GO:BP and REAC terms related to mitochondrial function and cellular respiration (**f**).

We performed a parallel analysis in excitatory neurons pooling the unique DEGs of organoid cluster 3 and 12 and compared this list to the human homologs of mouse DEGs from clusters 9, 11, and 15 (Fig. 3d). In this case, we found approximately twice the overlap that would be expected by chance among both increased and decreased genes (p < 1.4E-10). All significantly enriched GO:BP and Reactome (REAC) terms among down-regulated genes are shown (Fig. 3f) and all terms are related to cellular energy metabolism and mitochondrial function.

### Components of respiratory complexes II and IV are altered in mouse brain

The single-cell transcriptomic data from human 12-month organoids and mouse isocortex strongly indicated a long-term effect of the 3q29Del on mitochondrial function. More specifically, a top pathway dysregulated across 17 mouse clusters was *Mitochondrial Respiratory Chain Complex Assembly*. To test the hypothesis that the 3q29Del compromises the integrity of the mitochondrial respiratory chain at the protein level, we probed by Western blot for OXPHOS complex components in Percoll-isolated mitochondrial fractions from adult male and female mouse brains in control and 3q29Del backgrounds. Across 5 independent experiments with 2 mice per genotype pooled in each replicate, we observed selective decreases in components of Complexes II and IV (Fig. 4a-b, one sample Wilcoxon signed-rank test, p<0.05) indicating a shift in the stoichiometry of respiratory chain complexes.

**Fig 4.**
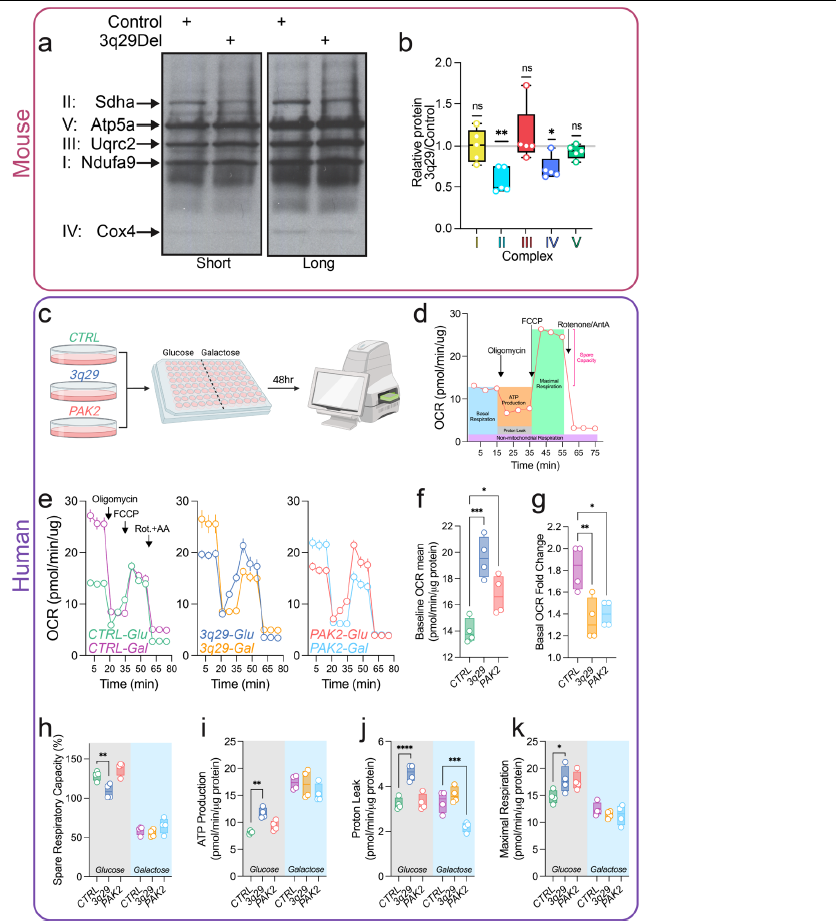
Mitochondrial phenotypes in 3q29 mice and engineered cell lines. Mitochondrial fractions from adult mouse brain lysates were found to have selective decreases in components of OXPHOS Complexes II and IV (**a**, quantified in **b**, N=5). Experimental overview for Seahorse assays and example calculations are shown in **c** and **d** respectively. Engineered HEK cells carrying a *TFRC-BDH1* hemizygous 3q29 deletion and cells completely lacking the 3q29-encoded gene *PAK2* were found to have elevated baseline oxygen consumption rate (OCR, **e**, **f**). *3q29* and *PAK2* cells displayed a blunted response to galactose medium (Gal, **e**, **g**). *3q29* cells had significantly reduced spare capacity (**f**), whereas spare capacity in *PAK2* cells was intact. *3q29* cells also exhibited significantly increased ATP production (**i**), proton leak (**j**), and maximal respiration (**k**) in baseline glucose conditions. Statistics: (**b**) one sample Wilcoxon signed-rank test (two-tailed), *p<0.05, **p<0.01; (**f**) one-way ANOVA main effect of genotype F(2, 9) = 17.24, p=0.0008, Dunnett multiple comparisons *CTRL vs. 3q29 ***p=0.0005, CTRL vs. PAK2 *p=0.0332*; (**g**) one-way ANOVA main effect of genotype F(2, 9) = 8.838, p=0.0075, Dunnett multiple comparisons *CTRL vs. 3q29 **p=0.0075, CTRL vs. PAK2 *p=0.0138*; (**h**) two-way ANOVA main effect of genotype F(2, 18) = 13.04 p=0.0003, Tukey comparison in glucose medium, *CTRL vs. 3q29 **p=0.0047, CTRL vs PAK2 p=0.2306*; (**i**) two-way ANOVA main effect of genotype F(2, 18) = 4.309 p=0.0296, Tukey comparison in glucose medium, *CTRL vs. 3q29 **p=0.0079, CTRL vs. PAK2 p=0.4812*; (**j**) two-way ANOVA main effect of genotype F(2, 18) = 31.16 p<0.0001 and treatment F(1, 18) = 21.45 p=0.0002, Tukey comparisons in glucose medium *CTRL vs. 3q29 ****p<0.0001, CTRL vs. PAK2 p=0.9503,* in galactose medium *CTRL vs. 3q29 p=0.4579, CTRL vs. PAK2 ***p=0.0007*; (**k**) two-way ANOVA main effect of treatment F(1, 18) = 57.31 p<0.0001, Sidak comparison in glucose medium *CTRL vs. 3q29 *p=0.0364, CTRL vs. PAK2 p=0.0572*. N=4 for all Seahorse experiments.

### Mitochondrial function is altered in 3q29Del engineered cell lines

Transcriptomic data from human 2-month 3q29Del organoids suggested an alteration in the timing and/or efficiency of the canonical glycolysis to OXPHOS shift in developing neural cells. We sought to mimic this shift by challenging engineered HEK-293T cell lines with galactose medium (Gal), which forces cells to utilize OXPHOS for energy production (Robinson et al., 1992). For these experiments, we used similar CRISPR/Cas9 methods to engineer a complete version of the hemizygous 3q29Del into HEK-293T cells (*TFRC-BDH1*) or to completely delete the 3q29 gene *PAK2* (Supp. Fig. 10), which was found to be among the most highly-expressed genes in the 3q29Del interval in nearly all mouse and human cell-types and was recently shown to be involved in cellular energy metabolism (Koranova et al., 2022). We then measured mitochondrial function in the Seahorse mitochondrial stress test, which isolates contributions of the respiratory chain complexes to oxygen consumption in cultured cells through sequential addition of inhibitor molecules. After 48hr of maintenance in galactose medium (Fig. 4c), *3q29* and *PAK2* cells did not up-regulate baseline mitochondrial function to the same degree of mock-edited *CTRL* cells (Fig 4e, g one-way ANOVA F(2, 9)=8.838, p=0.0075, Dunnett test *CTRL vs 3q29* **p=0.0075, *CTRL vs PAK2* *p=0.0138). Moreover, the baseline oxygen consumption rate (OCR) of *3q29* and *PAK2* cells in glucose-containing growth medium was significantly higher than in *CTRL* cells (Fig. 4e, f (genotypes split to improve visibility), one-way ANOVA main effect of group F(2, 9)=17.24, p=0.0008, Dunnett test *CTRL vs 3q29 \*\*\**p=0.0005, *CTRL vs PAK2* *p=0.0332).

Not all phenotypes were recapitulated by *PAK2* knockout. For example, *3q29* cells had a significant decrease in spare capacity (Fig. 4h two-way ANOVA main effects of genotype F(2, 18)=13.04, p=0.0003 and treatment F(1, 18) = 448.0, p<0.0001, *post-hoc* Tukey comparison in glucose medium *CTRL* vs *3q29* **p=0.0047), which was unaffected by loss of *PAK2* (p=0.2306). In fact, 3q29Del cells had almost no spare capacity, which may reflect increased vulnerability to metabolic challenges. In addition, only *3q29* cells had increased ATP production at baseline (Fig. 4i two-way ANOVA main effects of genotype F(2, 18)=4.309, p=0.0296 and treatment F(1, 18)=131.6, p<0.0001, *post-hoc* Tukey comparison *CTRL* vs *3q29* **p=0.0079) whereas *3q29* cells were found to have an increase in proton leak in glucose medium (Fig. 4j two-way ANOVA main effects of genotype F(2, 18)=31.16, p<0.0001 and treatment F(1, 18)=21.45, p=0.0002, *post-hoc* Tukey comparison *CTRL* vs *3q29* ****p<0.0001) but *PAK2* cells had significantly decreased proton leak in galactose medium (*CTRL* vs *PAK2* ***p=0.0007). Thus, *PAK2* is likely involved in the metabolic phenotypes of 3q29Del but is not solely sufficient.

Taken together, these experiments reveal that the 3q29Del produces a convergent mitochondrial phenotype at the level of transcriptome, protein expression and function.

## Discussion

As the strongest known genetic risk factor for SCZ, the 3q29Del is a high priority target for mechanistic investigation, which has thus far been limited. This study employed a novel cross-species strategy to identify transcriptomic phenotypes in human and mouse 3q29Del neural tissue and represents an important step toward understanding the neurobiological impact of this genetic variant. We leveraged the complementary advantages of two highly relevant experimental model systems to isolate the effects of the 3q29Del in the developing cortex and designed an analysis strategy to systematically filter widespread DEG findings down to the most likely processes and pathways for in-depth functional analysis. This approach led to neural mitochondria as a site of consistent gene dysregulation. Subsequent testing revealed changes in mitochondrial protein expression and function at the cellular level. These data strongly implicate the mitochondria as a site of impact for the 3q29Del. These results also highlight the strength of our cross-species approach, which rapidly and efficiently led to productive avenues for functional study.

In human cortical organoids, we found that *Oxidative Phosphorylation* was among the most frequently implicated pathways across cell-type clusters (Fig. 2). However, the direction of change was dependent on developmental timepoint: it was up-regulated in 2-month cells and decreased in 12-month cells. Moreover, the key glycolysis gene *LDHA* was found to be strongly decreased in multiple 2-month clusters that were overexpressing OXPHOS genes. These findings suggest that the glycolysis to OXPHOS transition, which is critical for neuronal differentiation and maturation (Zheng et al., 2016), may be disrupted in 3q29Del cells. We tested this prediction in an independent cellular model system, 3q29-engineered HEK cells, and found further evidence for a lack of metabolic flexibility; 3q29Del cells showed almost no spare capacity under baseline conditions and, when challenged with a galactose based medium to force aerobic respiration via OXPHOS, 3q29Del cells had a significantly blunted response (Fig. 4). In addition, ablation of the 3q29-encoded gene *PAK2* recapitulated two of these effects – decreased response to galactose medium and increased baseline aerobic respiration – while leaving spare capacity intact. These results suggest that *PAK2* is likely one of multiple 3q29 locus-encoded genes that contributes to metabolic phenotypes.

A major strength of the design of the current study is the utilization of human and mouse models as well as two time points *in vitro*. In both 12-month human cortical organoids and perinatal mouse cortical tissue, we found a widespread decrease in expression of genes related to mitochondrial energy production. In particular, down-regulated gene lists from 17 mouse clusters were enriched for *Mitochondrial Respiratory Chain Complex Assembly* (Fig. 2). In support of this transcriptomic prediction, we found evidence for a shift in the stoichiometry of respiratory chain complex proteins in mitochondrial fractions from mouse brain with specific decreases in protein components of complex II and IV (Fig. 4). This result indicates that the transcriptomic changes that we observed in perinatal mouse cortical tissue and 12-month *in vitro* human cortical organoids may translate into long-term, persistent effects at the protein level. Future work may be aimed determining if there is a mechanistic connection between the cellular energy metabolism phenotypes that we have described and the consistent finding that mice and human individuals with the 3q29Del are significantly smaller than expected (Baba et al., 2019; Rutkowski et al., 2019; Sanchez Russo et al., 2021).

Mitochondria have been previously implicated in the pathophysiology of neurodevelopmental CNV disorders and idiopathic schizophrenia (Ni et al., 2020; Rajasekaran et al., 2015). Interestingly, a CNV disorder with perhaps the most similar phenotypic profile to 3q29Del in human carriers, 22q11.2 deletion (22q11.2Del), harbors at least eight genes that encode mitochondria-linked proteins, several of which are also enriched at synapses (Maynard et al., 2008): *MRPL40, SLC25A1, PRODH, TXNRD2, AIFM3, COMT, RTL10, SNAP29* (Rath et al., 2021). Several mitochondrial phenotypes have now been reported in 22q11.2Del models. Similar to our findings, the activity of OXPHOS complexes I and IV was found to be decreased in human 22q11.2Del iPSC-derived neurons, which resulted in reduced ATP production (Li et al., 2019; Li et al., 2021). This suggests a potential convergent biology of mitochondrial dysfunction between 3q29Del and 22q11 deletion. This phenotype was attributed to haploinsufficiency of the 22q11.2Del locus gene *MRPL40*, which is a component of mitochondrial ribosome. Interestingly, loss of one copy of *Mrpl40* in mice is sufficient to produce short-term neuroplasticity phenotypes (Devaraju et al., 2017), potentially linking mitochondrial phenotypes to more well-established synaptic defects in SCZ models. A separate study that utilized a cross-species strategy to prioritize 22q11.2Del-associated effects in mouse brain and human patient fibroblasts identified the 22q11.2 gene encoding the mitochondrial citrate transporter *SLC25A1* as a key component of a dysregulated mitochondrial protein hub (Gokhale et al., 2019). Further studies indicated an interaction between SLC25A1 and MRPL40 at the protein level (Gokhale et al., 2021). Additionally, a large transcriptomic study of 22q11.2Del cortical organoids also found enrichment of DEGs related to mitochondrial function (Khan et al., 2020).

Other neurodevelopmental CNVs have been associated with mitochondrial phenotypes as well. The Williams Syndrome (deletion) and SCZ-associated (duplication) locus 7q11.23 contains the gene *DNAJC30*, which encodes a protein that interacts with the ATP synthase complex (Tebbenkamp et al., 2018). Complete loss of *DNAJC30* was found to disrupt sociability in mice and severely impair mitochondrial function in mouse neurons (Tebbenkamp et al., 2018). Human fibroblasts from individuals with Williams Syndrome (i.e. heterozygous for *DNAJC30*) were also found to have impaired mitochondrial function and reduced ATP production (Tebbenkamp et al., 2018). In addition, a recent study of reciprocal CNVs at the neurodevelopmental disorder associated locus 16p11.2 also found a strong signal for enrichment of DEGs related to energy metabolism and mitochondrial function in both mouse brain and cultured human neural cell lines (Tai et al., 2022). Another study of convergent biology in mouse models of the neurodevelopmental CNVs 1q21.1, 15q13.3, and 22q11.2 found dysregulation of a transcriptomic module related to neuronal energetics (Gordon et al., 2021). Finally, recent data indicates that complete loss of one of the top single gene risk factors for SCZ, *SETD1A* (Singh et al., 2022), impairs basal glycolysis and respiratory capacity in human neurons (Chong et al., 2022). Together, these data indicate that our findings in 3q29Del mouse and human cortical tissue fit with reports of other mitochondrial phenotypes associated with neurodevelopmental variants and suggest that neural mitochondria may be a key site of biological convergence downstream of these high-risk alleles.

Unlike 22q11.2, which encodes several proteins that function within mitochondria, the mechanistic link to 3q29 genes is not known. Our data implicates the highly expressed kinase *PAK2* in the metabolic phenotypes associated with 3q29Del, though likely in conjunction with additional driver genes, as predicted by our earlier network-based inferences on 3q29 neuropathology emerging upon loss of multiple functionally connected genes in the interval (Sefik et al., 2021). Mitochondria are involved in many cellular pathways and processes in addition to energy production including apoptosis signaling. Thus, our findings of transcriptomic and functional phenotypes at 3q29Del mitochondria fit with a previous report of increased susceptibility to apoptosis in *Drosophila* models (Singh et al., 2020). Further studies will be required to determine if mitochondrial phenotypes are a primary consequence of 3q29Del, and to identify the specific driver genes responsible for these effects.

Given the hierarchical structure of scRNA-Seq data, treating individual cells as independent sampling units can yield false positives in differential expression results due to the underestimation of true standard errors. To improve the reproducibility and validity of our findings, we focused our investigation exclusively on disrupted gene expression signals and corresponding signaling pathways with independent statistical support from two separate model systems. We note that while this approach increases our confidence in capturing true associations, additional signals relevant to disease mechanism may be hidden among unshared findings between model systems.

Bridging the gap between genetic risk and biological mechanisms is a major challenge for psychiatry. In this study, we sought to use systematic methods to identify the most salient, conserved transcriptomic effects of the SCZ-associated 3q29Del in disease-relevant tissues as an important step toward determining cellular and molecular phenotypes of this important variant. These findings should motivate further work to determine the mechanisms of these 3q29Del sequelae and their relevance to various clinical phenotypes.

**Supplemental Figure 1.**
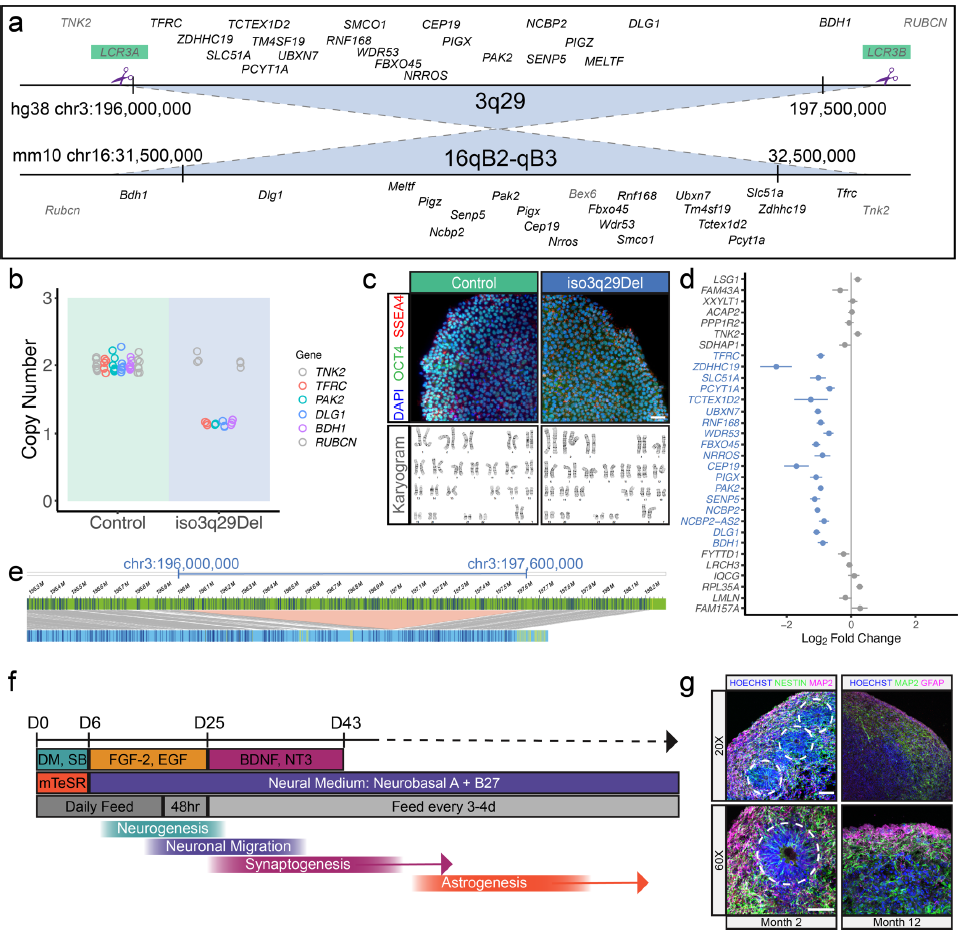
Isogenic human cortical organoids. The synteny of human and mouse 3q29Del loci is shown with purple scissors noting the approximate gRNA cut sites in the human LCRs (**a**). (**b**) TaqMan copy number assay data showing the loss of 1 copy of 3q29 genes *TFRC, PAK2, DLG1, BDH1* without affecting the copy number of the intact flanking genes *TNK* and *RUBCN*. (**c**) Isogenic 3q29Del cell lines retained normal cellular morphology (top) and normal karyotypes (bottom). (**d**) Expression of genes in the 3q29Del locus was decreased to approximately match gene copy number in isogenic 3q29Del iPSCs. (**e**) Bionano optical mapping showed no off-target structural variants aside from the 3q29Del (shown). (**f**) Schematic of differentiation protocol for dorsal forebrain cortical organoids (based on Sloan et al. 2018). (**g**) Example images of 2-month and 12-month cortical organoids with neural rosette-like structures highlighted with dashed line circles (scale bar = 50um).

**Supplemental Figure 2.**
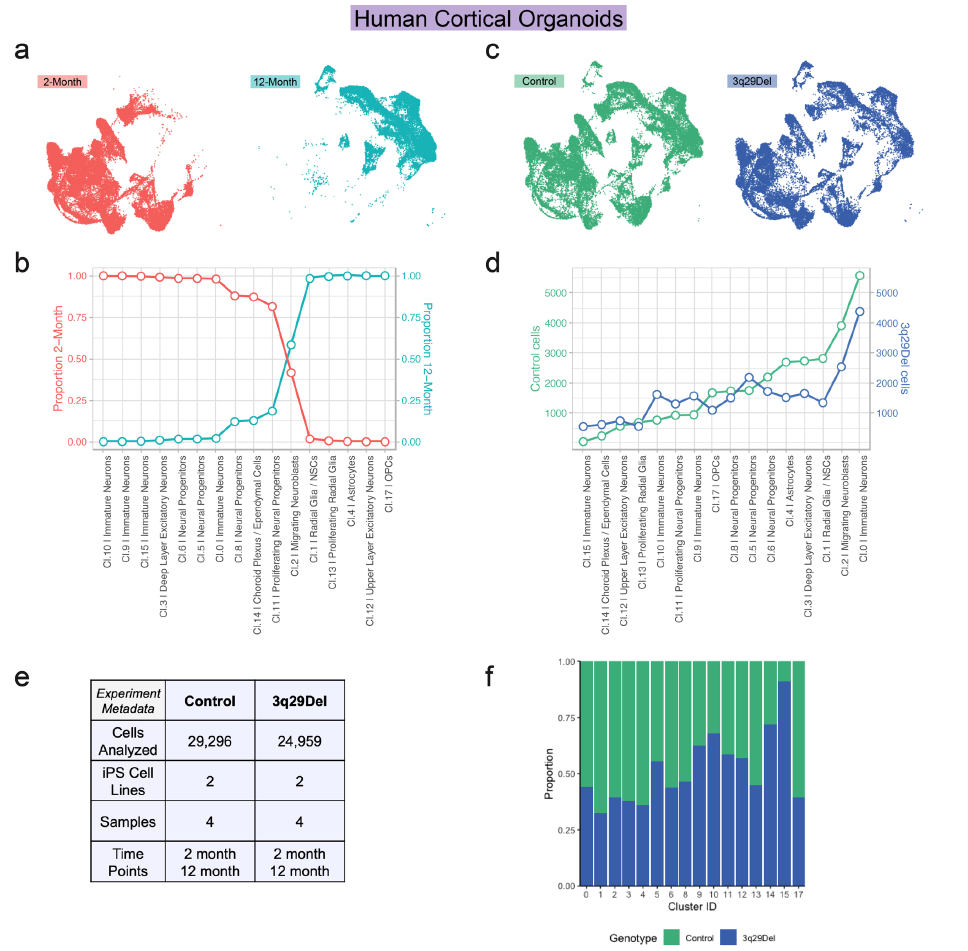
Human cortical organoid cluster proportions by time point and genotype. (**a**) Human cortical organoid UMAP projections split by time point. (**b**) Proportion of cells in each organoid cluster from each time point. (**c**) Human cortical organoid UMAP projections split by genotype. Number (**d**) and proportion (**f**) of cells in each organoid cluster from each genotype. (**e**) Human experiment metadata. Abbreviations: OPC, oligodendrocyte progenitor cells; NSC, neural stem cells; cl, cluster.

**Supplemental Figure 3.**
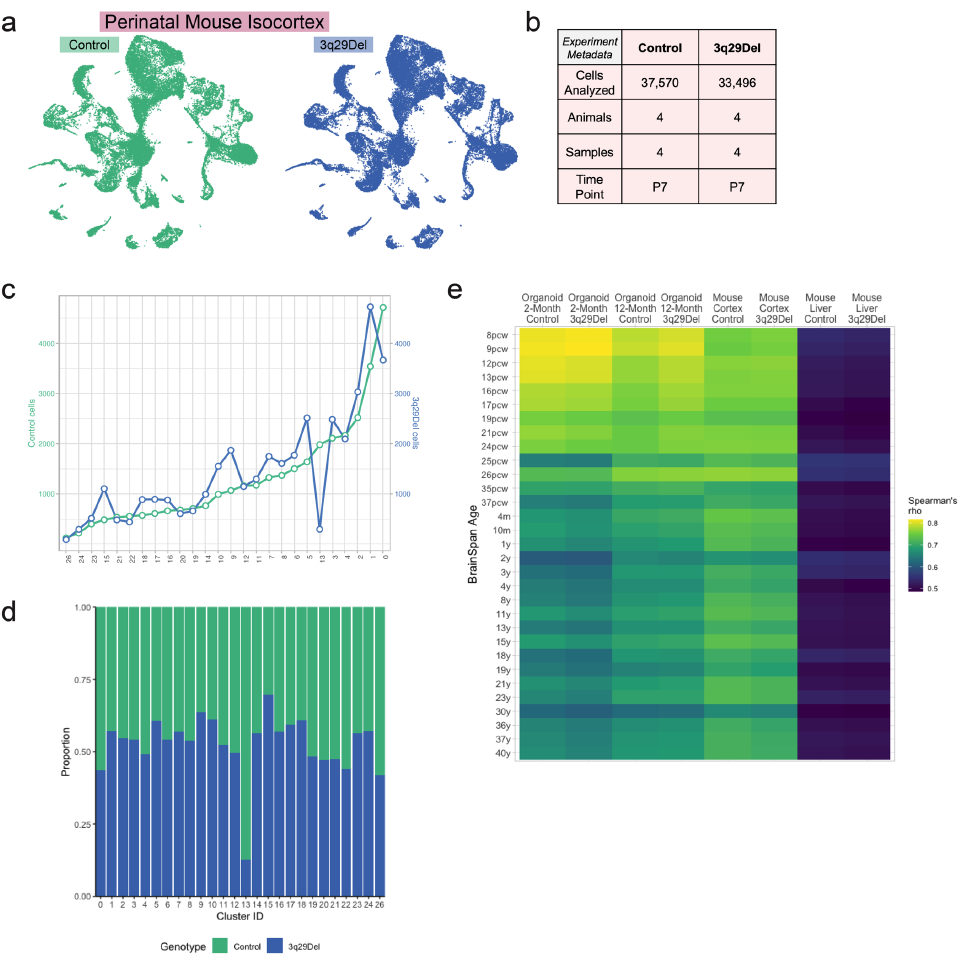
Mouse isocortex cluster proportions by genotype. (**a**) Mouse isocortex UMAP projections split by genotype. (**b**) Mouse experiment metadata. Number (**c**) and proportion (**d**) of cells in each mouse cluster from each genotype. (**e**) Spearman correlations of the average gene expression profiles of human cortical organoids, human homologs of mouse isocortex, and human homologs of adult mouse liver samples (from Pollak et al. 2022) compared to BrainSpan gene expression data from post-mortem human brains (Kang et al., 2011). Abbreviations: pcw, post-conception weeks (prenatal); m, months (postnatal); y, years (postnatal).

**Supplemental Figure 4.**
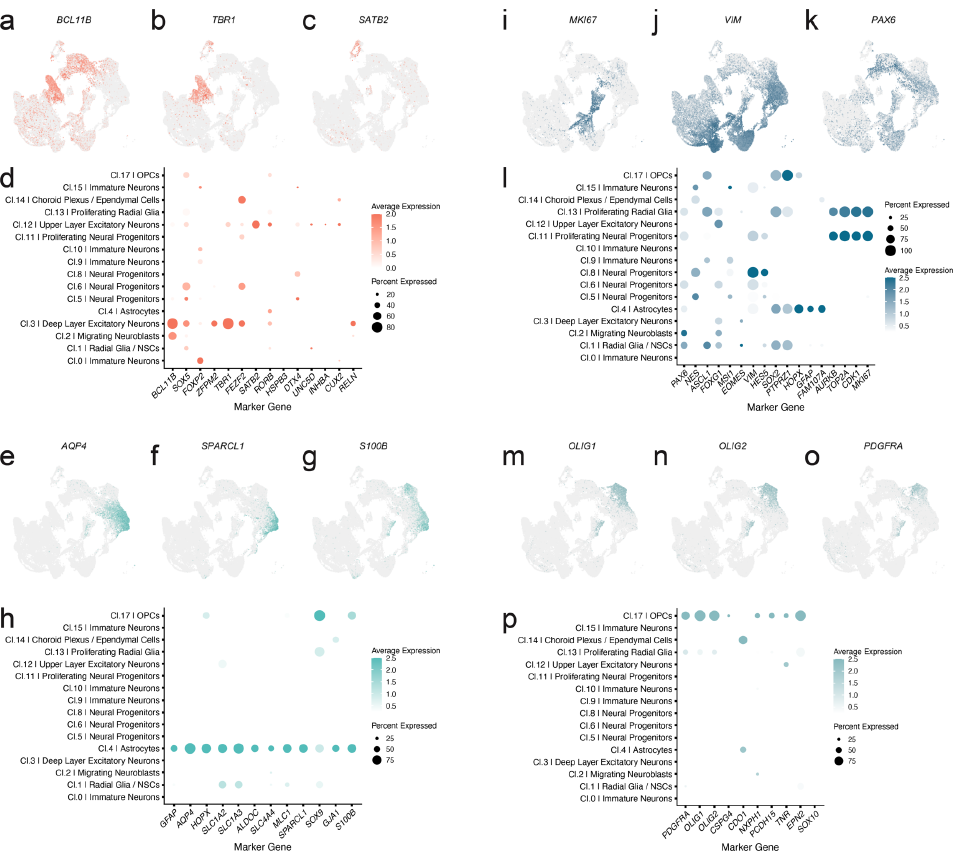
Human organoid cell-type marker gene expression. Clusters were annotated for cell-type based on differential marker gene expression, as curated from the literature. Mature excitatory neurons (**a**-**d**) were defined by expression of deep layer (*BCL11B*, *TBR1*) or upper layer (*SATB2*, *CUX2*) markers. Neural progenitors and radial glia/neural stem cells (NSCs) (**i**-**l**) were abundant in cortical organoids. Proliferating progenitors and radial glia/NSCs expressed *MKI67*. Astrocytes (**e**-**h**) were only found in 12-month cells and expressed *GFAP*, *AQP4*, and *HOPX*. Oligodendrocyte progenitor cells (OPCs, **m**-**p**) expressed established marker genes such as *PDGFRA*, *OLIG1*, and *OLIG2*. Dot size in dot plots displays the fraction of cells expressing representative marker genes in individual clusters (minimum 10%) and dot color represents the scaled average expression levels. Example UMAP visualizations are provided in conjunction with dot plots to display the distribution of major cell populations expressing well-known marker genes relative to non-expressing cells.

**Supplemental Figure 5.**
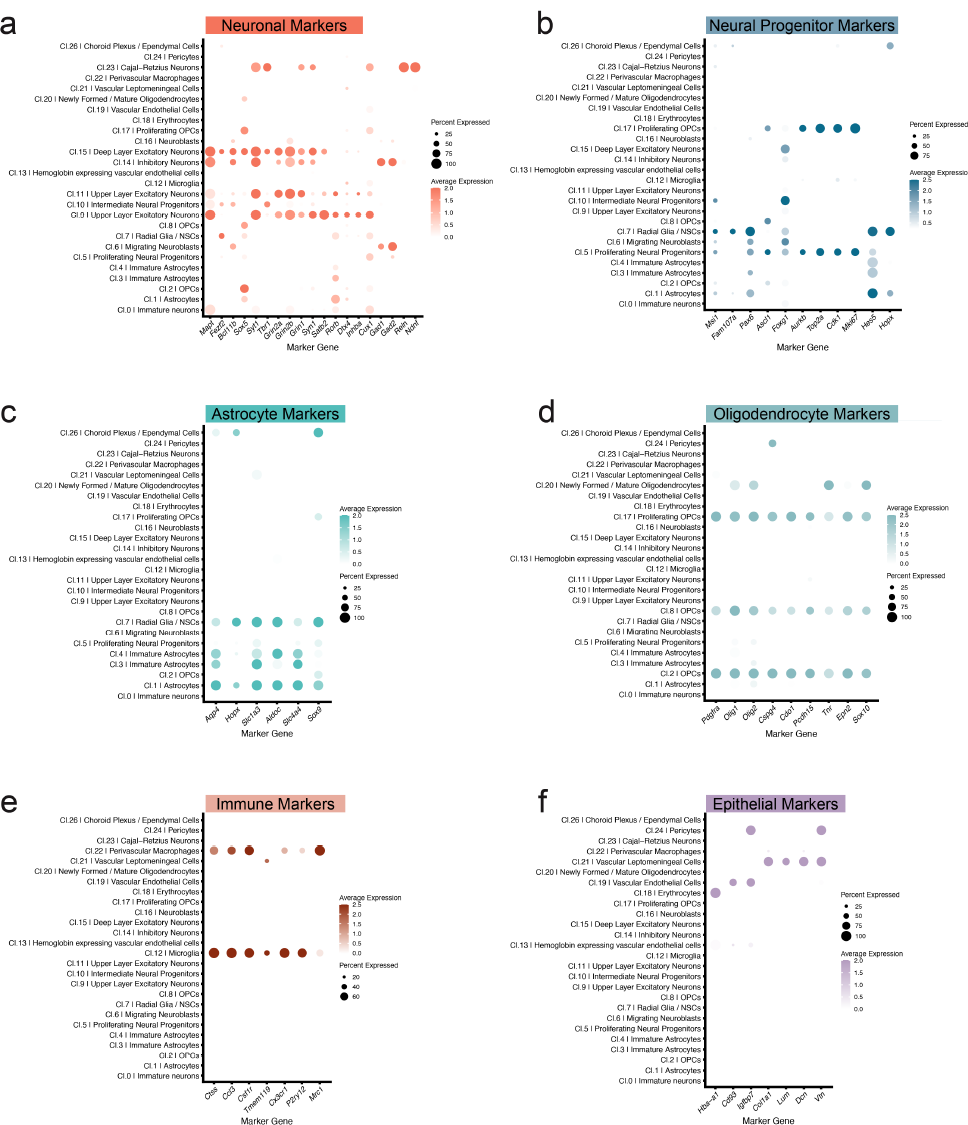
Mouse cell-type marker gene expression. Five clusters of neurons were identified in mouse isocortex (**a**): *Satb2* and *Cux1* expressing upper layer excitatory neurons (cl. 9 and 11), *Gad1* and *Gad2* expressing inhibitory neurons (cl. 14), *Bcl11b* and *Tbr1* positive deep layer excitatory neurons (cl. 15), and *Reln* expressing layer I Cajal-Retzius neurons (cl. 23). All neuron clusters express the well-known pan-neuronal marker *Syt1*. Markers for proliferating neural progenitors (cl. 5) and radial glia/neural stem cells (NSCs, cl. 7) are shown in **b**. One cluster (cl. 1) contained astrocytes positive for canonical markers such as *Aqp4* and *Sox9* though others (cl. 3-4) also expressed markers at lower levels (obscured by plot scaling). Three clusters of oligodendrocyte progenitor cells (**d**, OPCs, cl. 2, 8, 17) were identified along with newly-formed / mature oligodendrocytes (cl. 20). Microglia expressing *Cx3cr1* and *P2ry12*, macrophages expressing *Mrc1* (**e**), and other non-neuronal cell-types such as vascular, leptomeningeal, endothelial and ependymal cells (**f**) were also present. Dot size in dot plots displays the fraction of cells expressing representative marker genes in individual clusters (minimum 10%) and dot color represents the scaled average expression levels.

**Supplemental Figure 6.**
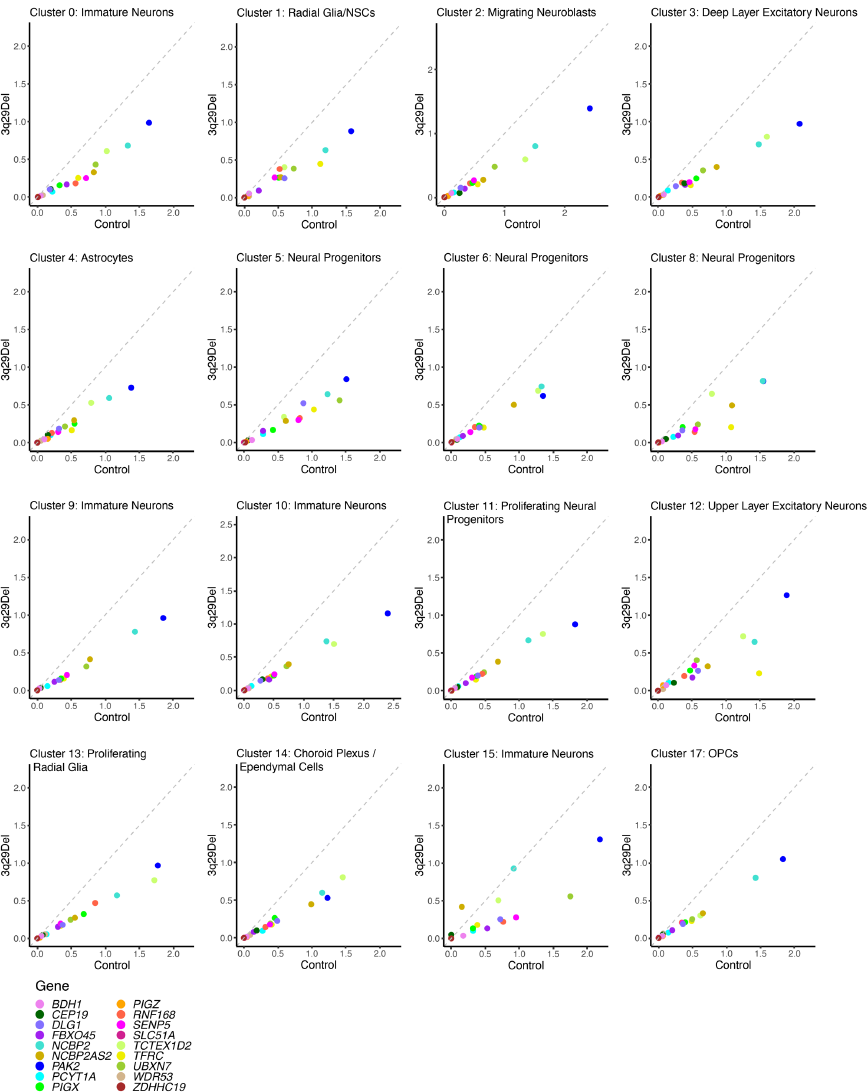
Relative expression of 3q29 genes in human cortical organoids. The average expression profile of genes in the 3q29Del locus are plotted, 3q29Del (y-axis) relative to Control (x-axis). In nearly every cluster, expression of all 3q29 genes is higher in Control cells. The dashed diagonal line indicates equal average expression between genotypes (x=y). Abbreviations: OPC, oligodendrocyte progenitor cells; NSC, neural stem cells.

**Supplemental Figure 7.**
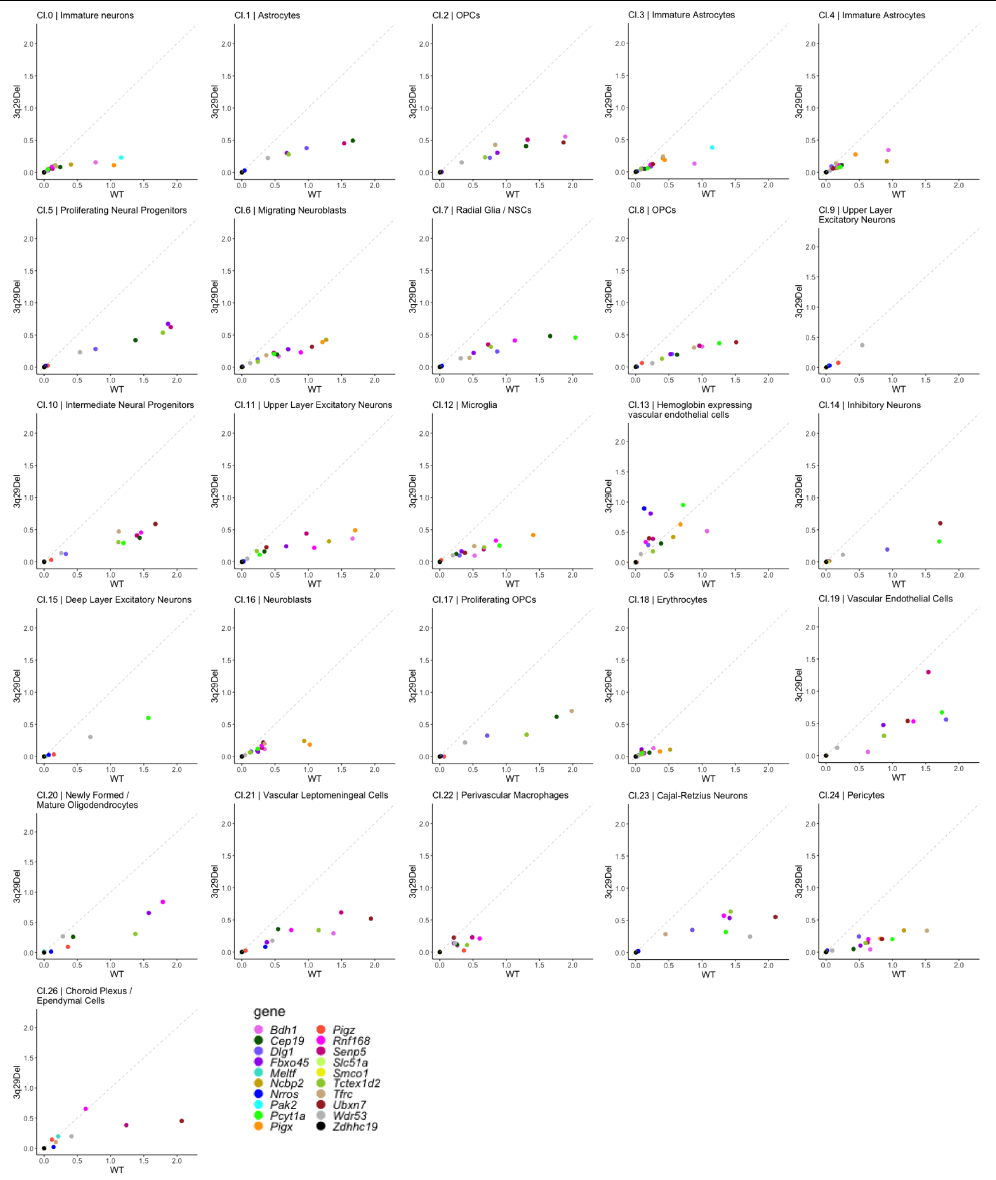
Relative expression of 3q29 genes in P7 mouse isocortex. The average expression profile of genes in the mouse homologous 3q29Del locus are plotted, 3q29Del (y-axis) relative to Control (WT, x-axis). In nearly every cluster, expression of all 3q29 genes is higher in Control cells. The dashed diagonal line indicates equal average expression between genotypes (x=y). Abbreviations: OPC, oligodendrocyte progenitor cells; NSC, neural stem cells.

**Supplemental Figure 8.**
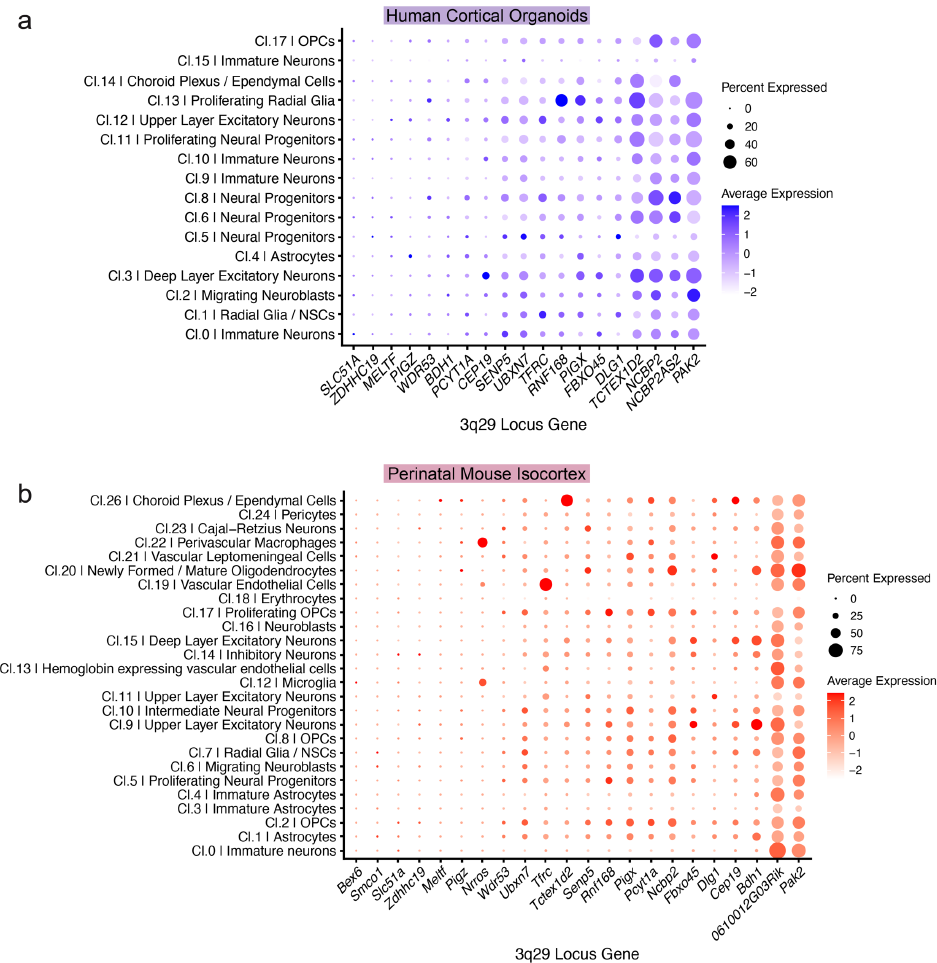
Relative expression of 3q29 genes across cortical cell-types. Dot plots visualizing the relative expression of 3q29 locus transcripts across clusters in human cortical organoids (**a**) and the expression of mouse homologs of human 3q29 locus genes in mouse cortex clusters (**b**), ordered by approximate total expression. Dot size displays the fraction of cells expressing 3q29 genes in individual clusters and dot color represents the scaled average expression levels. 3q29 interval genes that were not detected in human organoids: *SMCO1, NRROS, TM4SF19*. 3q29 interval homolog not detected in mouse cortex: *Tm4sf19*. Abbreviations: OPC, oligodendrocyte progenitor cells; NSC, neural stem cells; cl, cluster.

**Supplemental Figure 9.**
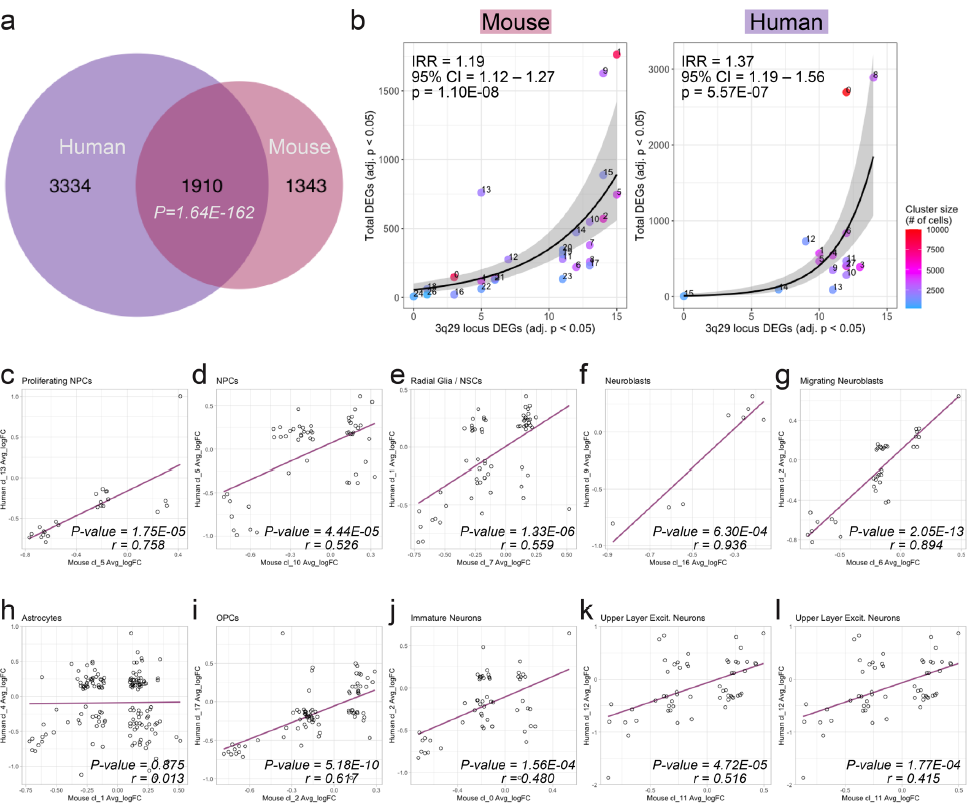
Mouse – human comparison. In (a) more than half of the human homologs of all unique mouse DEGs from any cluster were also found to be differentially expressed in at least one human organoid cluster. The total number of DEGs in a given mouse or human cluster is not significantly related to cluster size (number of cells in a cluster) but to the number of 3q29 locus genes that were differentially expressed (negative binomial regression, p<0.0001). Estimated rate ratios for total number of DEGs are indicated above each plot for a unit increase in the number of differentially expressed 3q29 locus genes found per cluster, while keeping cluster size constant. (b). (c-l) Scatterplots showing the relationship between average log fold change (Avg_logFC) values of shared DEGs in representative clusters of mouse vs human cell-types. P-values and correlation coefficients from Pearson correlations are indicated above each plot. Purple lines represent the line of best fit. Abbreviations: DEG; differentially expressed gene; IRR, incidence rate ratio; CI, confidence interval; OPC, oligodendrocyte progenitor cells; NSC, neural stem cells; NPC, neural progenitor cells; Excit. excitatory; cl, cluster.

**Supplemental Figure 10.**
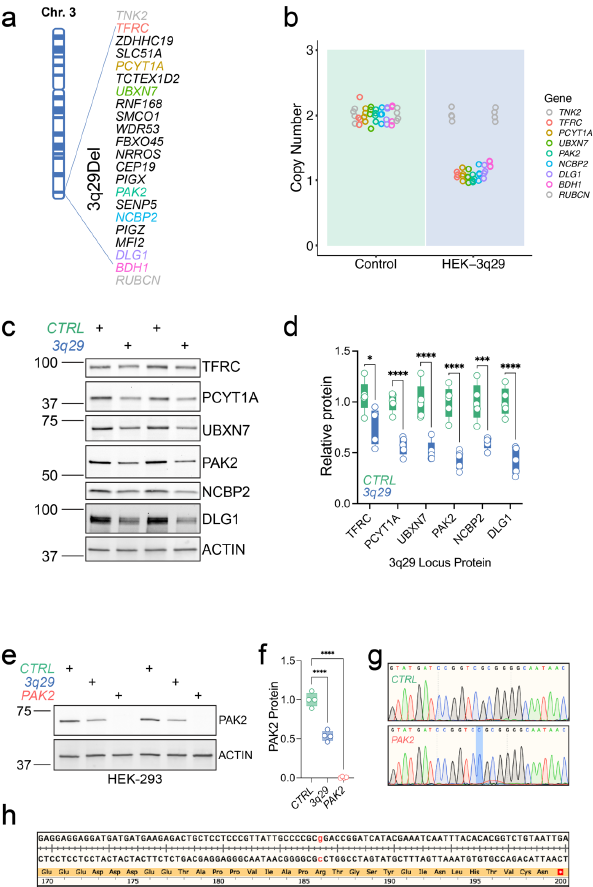
Engineered HEK cell lines. The protein-coding genes of the 3q29 locus are shown (**a**) with those that would be probed by Western blot highlighted in blue. TaqMan Copy Number assays for 3q29 deletion locus genes *TFRC, PAK2,* and *DLG1* showed reductions consistent with hemizygosity whereas the most distal 3q29 deletion gene *BDH1* was not clearly heterozygous or unaffected. Western blots (**c**) from whole cell lysates showed clear and significant reductions in all 3q29-encoded proteins tested (**d**). BDH1 was not reliably detected in these cell lysates. Western blot analysis (**e**-**f**) of whole cell lysates from *3q29* and *PAK2* cells compared to *CTRL* showed the expected 50% and complete depletion of PAK2 protein. Sanger sequencing of PCR-amplified gDNA from *PAK2* cells showed an insertion (**g**, highlighted in blue) that is predicted to produce a new premature stop codon (**h**, insertion highlighted in red). The PAK2 antibody used in **c** and **f** is directed against an N-terminal fragment and no lower molecular weight bands were detected, which indicates that no PAK2, even in truncated form, is produced in these cells.

## Acknowledgments

This study was supported by NIMH F32 MH124273 (RHP), a NARSAD Young Investigator Grant from the Brain & Behavior Research Foundation (RHP), NIMH R56 MH116994 (DW, STW, JGM), R01 MH110701 (GJB and JGM), R01 MH118534 (JGM). Research reported here was also supported in part by Imagine, Innovate and Impact (I3) from the Emory School of Medicine, a gift from Woodruff Fund Inc., and through the Georgia CTSA NIH award (UL1-TR002378) and by the University Research Committee of Emory University. This study was also supported in part by the Emory Integrated Genomics Core (EIGC), Emory Integrated Computational Core (EICC), Emory Integrated Cellular Imaging, Emory Flow Cytometry Core, and the Emory Stem Cell Core, which are subsidized by the Emory University School of Medicine and are part of the Emory Integrated Core Facilities.

